# Unsupervised correction of gene-independent cell responses to CRISPR-Cas9 targeting

**DOI:** 10.1101/228189

**Authors:** Francesco Iorio, Fiona M Behan, Emanuel Gonçalves, Shriram G Bhosle, Elisabeth Chen, Rebecca Shepherd, Charlotte Beaver, Rizwan Ansari, Rachel Pooley, Piers Wilkinson, Sarah Harper, Adam P Butler, Euan A Stronach, Julio Saez-Rodriguez, Kosuke Yusa, Mathew J Garnett

**Affiliations:** European Molecular Biology Laboratory – European Bioinformatics Institute, Cambridge – UK; Wellcome Sanger Institute, Cambridge – UK; GlaxoSmithKline, Stevenage – UK; RWTH Aachen University, Faculty of Medicine, Joint Research Center for Computational Biomedicine Aachen – Germany; Open Targets, Cambridge – UK; Heidelberg University, Faculty of Medicine, Institute for Computational Biomedicine, Bioquant, Heidelberg, Germany

**Keywords:** CRISPR-Cas9, Genetic Screens, Cancer, Gene copy number, bias correction

## Abstract

**Background:** Genome editing by CRISPR-Cas9 technology allows large-scale screening of gene essentiality in cancer. A confounding factor when interpreting CRISPR-Cas9 screens is the high false-positive rate in detecting essential genes within copy number amplified regions of the genome. We have developed the computational tool *CRISPRcleanR* which is capable of identifying and correcting gene-independent responses to CRISPR-Cas9 targeting. CRISPRcleanR uses an unsupervised approach based on the segmentation of single-guide RNA fold change values across the genome, without making any assumption about the copy number status of the targeted genes.

**Results** Applying our method to existing and newly generated genome-wide essentiality profiles from 15 cancer cell lines, we demonstrate that CRISPRcleanR reduces false positives when calling essential genes, correcting biases within and outside of amplified regions, while maintaining true positive rates. Established cancer dependencies and essentiality signals of amplified cancer driver genes are detectable post-correction. CRISPRcleanR reports sgRNA fold changes and normalised read counts, is therefore compatible with downstream analysis tools, and works with multiple sgRNA libraries.

**Conclusions** CRISPRcleanR is a versatile open-source tool for the analysis of CRISPR-Cas9 knockout screens to identify essential genes.

## Background

CRISPR-Cas9-based genome editing techniques are transforming the landscape of genetic studies [1,2]. The high efficiency and specificity of the CRISPR-Cas9 system to mutagenise genes through the introduction of DNA double strand breaks (DSB), either at the level of individual genes or at genome-wide scale, enables the systematic investigation of loss-of-function phenotypes.

We and others have developed genome-wide pooled CRISPR knock-out (CRISPR-KO) screening strategies [3–5]. A prominent application of CRISPR-KO screens is the systematic identification of genes that are essential for cancer cell fitness to identify strategies for the development of novel targeted therapies. These studies typically introduce Cas9 endonuclease into cells, followed by or alongside the introduction of a library of pooled sgRNAs targeting the genome. The library usually contains multiple single guide RNA (sgRNA) targeting each gene to facilitate a robust identification of essential genes. Analysis strategies compare the abundance of sgRNAs between control and test samples to determine which sgRNAs are differentially under-represented, thus targeting a gene that is potentially essential to the fitness of the cancer cells. Several groups have performed these types of screens to identify novel drug targets [6,7]. A recent landmark study has reported gene essentialities in 342 cancer cell lines [8]. This will empower association studies between gene essentialities and genomic/transcriptomic features to develop biomarkers for patient stratification.

One drawback of the CRISPR-KO screening system is caused by its mode of action, namely DSB induction. DSBs trigger a DNA damage response which can cause cell cycle arrest and in some cases cell death [9–11]. This is problematic when performing whole-genome CRISPR-KO screens in cancer cells because of frequent copy number (CN) alterations in their genome, resulting in widespread Cas9 induced DNA damage. Consequently, DSBs at genes in amplified regions result in depletion of these genes in a pooled CRISPR-KO screen regardless of their essentiality, and thus they are erroneously called as fitness genes. This can result in a high false-positive rate and correcting for this CN-associated effect is crucial for the interpretation of CRISPR-KO screening results. Solutions proposed thus far encompass scanning the dataset for biased regions and their removal from downstream analysis [12], resulting in the exclusion of potentially biologically relevant genes residing in CN-amplified regions, or to apply a piecewise linear model to infer true gene dependencies based on CN profiles across large panels of cell lines [8].

During the analysis of CRISPR-KO data we identified a number of instances for which existing approaches for correcting bias in CRISPR-KO data were unsuitable or hampered further downstream analyses. To address this, we developed *CRISPRcleanR*, a computational approach implemented in open-source R and a Python packages, which identifies biased genomic regions from CRISPR-KO screens in an unsupervised manner and provides both corrected read count and log fold change (logFC) values of individual sgRNAs in such regions. Our method reduces false positive calls while keeping the true positive rate of known essential genes largely unchanged, and allows the detection of essential genes even within focally amplified regions.

## Results

### Gene-independent responses in CRISPR-KO screens

We performed genome-wide CRISPR-KO screens on 15 human cancer cell lines (hereafter called ‘Project Score’), which are a subset of the Genomics of Drug Sensitivity in Cancer (GDSC) collection (**Supplementary Table S1**) [13,14]. This involved six tumour types with different mutational processes, including high frequency of single-nucleotide variants (large intestine, lung, and melanoma) and CN variation (breast and ovary). We used the Sanger Institute CRISPR library (version 1.0) targeting 18,010 genes (90,709 sgRNAs; ∼5 sgRNAs per gene) [6]. The screens showed high consistency between technical replicates in each cell line (median average correlation for sgRNA counts = 0.83) and readily discriminated between pre-defined fitness essential (FE) and non-essential genes (median area under the Receiver Operating Characteristic curve (AUROC) = 0.92) (**Supplementary Fig. S1**) [15]. Additionally, a high true positive rate (TPR, or recall) was observed for known essential genes assembled from the Molecular Signature Database (MsigDB) [16] and from literature [17] (median TPR across gene sets and cell lines = 85% at 5% FDR).

When comparing CRISPR data and CN profiles for each line, we confirmed a large negative effect for logFCs of sgRNAs targeting genes in CN-amplified regions, particularly with CN ≥ 8 (**Supplementary Fig. S2** and **Supplementary Table S2**). Notably, sgRNA targeting CN-amplified (CN ≥ 8) non-expressed genes (FPKM < 0.05) were significantly more depleted in six cell lines than the rest of the sgRNA in the whole library. For three cell lines (HT55, EPLC-272H, and MDA-MB-415), the negative effect on logFC of sgRNA in CN-amplified regions was comparable or greater than for FE genes (**Supplementary Fig. S3** and **Supplementary Table S2**). Collectively, using independent data, our analysis confirms the systematic negative bias on sgRNA logFC values in particular regions of the genome, which are enriched for CN amplifications.

### Variable effect of amplification on responses to CRISPR-Cas9 targeting

To gain greater insight into CN-associated biases, we performed a detailed analysis of the relationship between sgRNA logFC values and CN at the level of individual CN segments (**Fig. 1a** and **Supplementary Fig. S4**). For some cell lines, the negative bias on average logFC values within segments was positively correlated with CN values (EPLC-272H, NCI-H520, OVCAR-8, TOV-21G and SW48). In other cell lines the bias effect on average logFC was not observed (MDA-MB-436), plateaued (NCI-H2170), or fluctuated as CN varied (MDA-MB-453, HT55 and HuP-T3). These effects were preserved when only considering sgRNA targeting non-expressed genes (**Fig. 1b** and **Supplementary Fig. S4**), demonstrating that the negative logFCs are most likely independent of true gene essentiality. In addition, we observed a wide range of average logFC values for segments of a given CN (**Fig. 1a, b**), and this is often larger than the variation between segments of different CN, indicating that CN alone does not capture all of the observed bias variance.

**Fig. 1:**
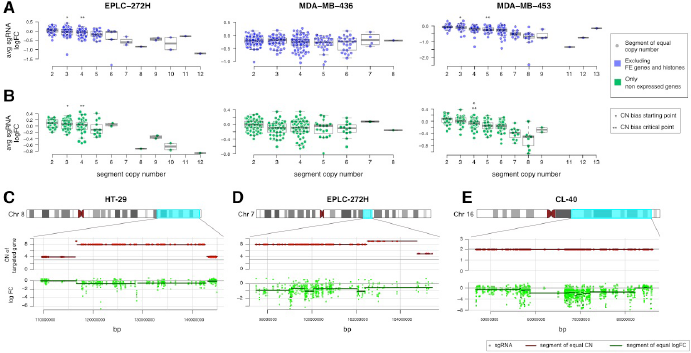
Heterogeneous gene-independent responses to CRISPR-Cas9 targeting. (A) Average logFC values of sgRNA within segments of equal CN (excluding FE and histones) for three cell lines. Each circle represents a CN segment of the indicated copy number. Asterisks mark the CN at which a significance difference (Welchs t-test, p < 0.05) is initially (starting point) and continuously (critical point) observed compared to logFC values at CN = 2. Box-plots show the median, inter-quartile ranges and 95% confidence intervals. (B) Same as for A but considering only non-expressed genes (FPKM < 0.05). (C, D, E) Segments of equal gene copy number and segments of equal sgRNA logFCs for selected chromosomes in three cell lines.

Furthermore, although in the majority of instances CN segments matched segments of equal sgRNA logFCs (**Fig. 1c**), we identified several CN segments with discontinuous logFC patterns (**Fig. 1d**). Additionally, regions of consistently depleted sgRNAs were identified also in diploid regions of the genome. For example, the cell line CL-40 harbours two copies of chromosome 16, but several contiguous genes (of which many are not expressed) in region 16q23 exhibited a negative logFC across targeting guides (**Fig. 1e**).

Our results indicate that biases observed in CRISPR-KO screens are often associated with CN alterations but are heterogeneous, with poorly understood variation between segments of differing CN, and variation within segments of the same CN. Taken together, these results highlights the value of an unsupervised approach, not dependent on CN alone, to correct for biased regions in CRISPRKO data.

### CRISPRcleanR corrects bias in CRISPR-Cas9 datasets

In order to detect biased regions in an unsupervised manner and correct corresponding sgRNA logFCs in CRISPR-KO screening data, we developed *CRISPRcleanR*, a computational approach implemented in open-source R and Python packages. CRISPRcleanR applies a circular binary segmentation algorithm, originally developed for array-based comparative genomic hybridization assay [18,19], directly to the genome-wide patterns of sgRNA logFCs across individual chromosomes in a cell line. It then detects genomic segments containing multiple sgRNAs with sufficiently equal logFCs. If these segments contain sgRNAs targeting a minimum number of distinct genes then the sgRNA in the segment are most likely responding to CRISPR-Cas9 targeting in a gene-independent manner, and logFCs values are corrected via mean-centering. Median-based centering can also be applied for experimentally variable data or in the presence of many outliers.

CRISPRcleanR embeds functions from the *DNAcopy* R package [20] allowing users to customise their arguments. Furthermore, it has several features that make it statistically robust, versatile and practical for downstream applications: (i) it works in an unsupervised manner, requiring no chromosomal CN information nor *a priori* defined sets of essential genes; (ii) it implements a logFC correction, making depletion scores for all genes usable in follow up analyses; (iii) it examines logFC at the sgRNA level to gain resolution and to account for different levels of sgRNA on-target efficiency, and enables the subsequent use of algorithms to call gene depletion significance that require input data at the sgRNA level (e.g. BAGEL [21]); (iv) by applying an inverse transformation to corrected sgRNA logFCs, it computes corrected sgRNA counts, which are required as input for commonly used mean-variance modeling approaches, such as MAGeCK [22], to call gene depletion/enrichment significance; (v) lastly, CRISPRcleanR corrects logFC values using data from an individual cell line and with invariant performances, unlike other computational correction approaches whose performances depend on the number of analysed cell lines [8]; as a consequence, CRISPRcleanR is suitable for the analysis of data from both small- and large-scale CRISPR-KO studies.

When applied to Project Score data, CRISPRcleanR effectively corrected the bias in sgRNA logFCs over a wide range of chromosomal segments with variable CN alterations. Furthermore, this included detection and correction of different level of biases in sgRNA logFCs within an individual segment of equal CN (**Fig. 2a, b**). An immediate result of the application of CRISPRcleanR to our data was that biases in particularly high CN regions were strongly attenuated over all the cell lines (**Fig. 2c**).

**Fig. 2:**
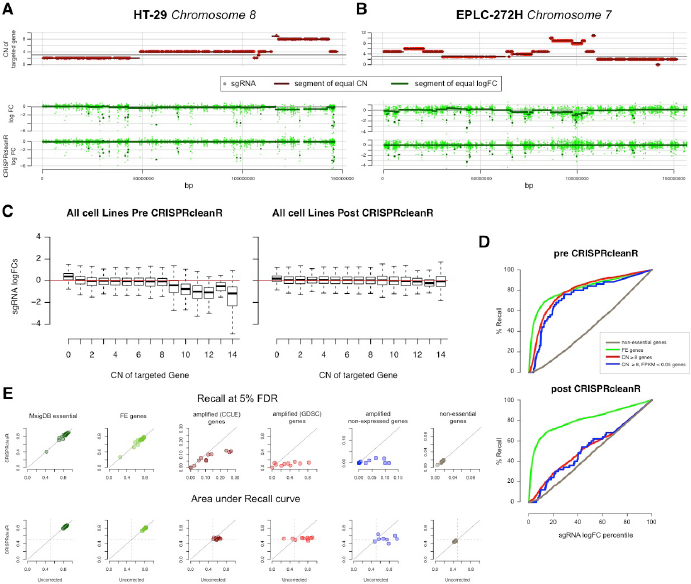
Unsupervised detection of segments of equal sgRNA logFCs and their correction. (A and B) Example segments of equal gene copy number and equal sgRNA logFC values detected and corrected by CRISPRcleanR in two cell lines. (C) logFC values of sgRNAs of the entire library for all cell lines grouped according to the copy number of their targeted gene before (left) and after (right) CRISPRcleanR correction. Box-plots show the median, inter-quartile ranges and 95% confidence intervals. (D) Recall curves of sgRNA when classified as targeting amplified genes, amplified non-expressed genes, FE genes, and non-essential genes before and after CRISPRcleanR correction, for an example cell line (EPLC−272H). (E) Assessment of CRISPRcleanR correction comparing Recall at 5% FDR (top row) or area under the Recall curve (AURC, bottom row) of genes in six predefined gene sets based on their uncorrected or corrected logFCs (averaged across targeting sgRNAs).

Overall, CRISPRcleanR reduced the recall of sgRNAs targeting CN-amplified regions, including sgRNAs targeting CN-amplified non-expressed genes, towards expectation when classifying the whole library of sgRNAs based on their logFCs (**Fig. 2d**). The correction was also consistently observed at the gene level (average logFCs of targeting sgRNAs) across all screened cell lines at a fixed 5% FDR, with a median reduction in recall equal to 72% and 88%, respectively for CN-amplified and CN-amplified non-expressed genes (**Fig. 2e** and **Supplementary Table S3**). This reduction was also observed at the level of the area under the overall recall curves (AURCs), thus independent of a fixed depletion significance threshold. Specifically, we observed the median AURCs across all cell lines shifting from 0.74 to 0.51 (*p* = 0.02, Welch’s two sample t-test) and from 0.7 to 0.5 (*p* = 0.01), respectively, for CN-amplified and CN-amplified non-expressed genes (**Fig. 2e** and **Supplementary Table S3**). The reduction in AURC was independent of whether amplified genes in cell lines were identified using CN data from the GDSC or the cancer cell line encyclopedia (CCLE). In contrast, for the MsigDB known essential genes and the FE genes, the reduction was negligible at less than 2%, with median AURCs preserved at ≥ 0.82.

Excluding from the essentiality profiles the sgRNAs targeting *a priori* known essential genes (taken from MSigDB) before CRISPRcleanR correction yielded very similar results as when imposing the constraint that, for a segment to be corrected, it must contain sgRNA targeting *n* = 3 different genes (**Supplementary Fig. S5**). This was determined by performing several correction attempts varying *n* and considering or not FE and other MSigDB essential genes. Thus, CRISPRcleanR can be used in a completely unsupervised setting, without making any assumption on gene essentiality.

### CRISPRcleanR is effective using multiple sgRNA libraries

To investigate the versatility of CRISPRcleanR we assessed its performance across different libraries of sgRNAs. For the purpose of comparability we initially used our previously published dataset derived from screening the HT-29 cell line with the Brunello [23] and Whitehead [12] libraries, using the same lentiviral vector as our library [24]. Of note, despite all three libraries targeting 17,646 overlapping genes, fewer than 5% of the 19-mer gRNA in the libraries are overlapping in sequence. A similar reductions in recall for CN-amplified genes (mean = 40 ± 2.7 %), CN amplified non-expressed genes (45 ± 5.7 %), fitness essential genes (2 ± 0.47 %), and non-essential genes (mean = −3.8 ± 1.81 %) was observed across all three libraries (**Fig. 3a, b**). As a specific example, all three libraries showed matching patterns of biased logFCs in the same CN-amplified genomic region spanning the proto-oncogene *MYC* on chromosome 8 (**Fig. 3c**). CRISPRcleanR corrected the sgRNA logFC values for this bias in all three libraries.

**Fig. 3:**
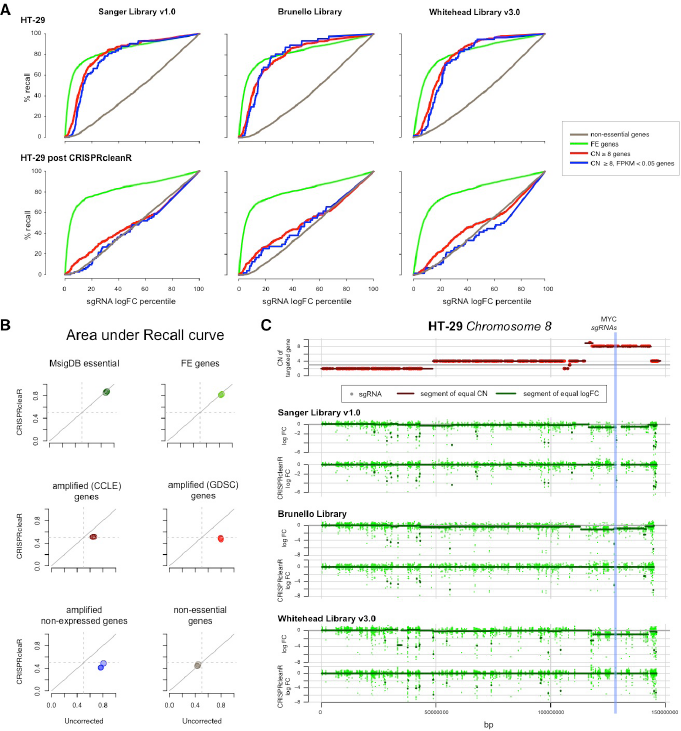
CRISPRcleanR is effective with multiple different sgRNA libraries. (A) Recall curves for three sgRNA libraries when classifying sgRNAs targeting amplified genes, amplified non-expressed genes, FE genes, and non-essential genes using sgRNA logFCs before (first row of plots) and after (second row of plots) CRISPRcleanR correction. (B) Variation of the area under the recall curve for sgRNAs targeting genes in six predefined sets, based on their uncorrected/corrected logFCs, across the three different libraries (one circle per library). (C) Segments within chromosome 8 of equal gene copy number juxtaposed to segments of equal sgRNA logFCs before and after CRISPRcleanR in HT-29 cells screened with three different sgRNA libraries. The position of *MYC* is shown with a blue line.

To further evaluate the compatibility of CRISPRcleanR with different sgRNA libraries, we tested it on an independent dataset of 342 cell lines using the Avana library from Project Achilles [8] (**Supplementary Fig. S6**). We observed a reduction of false positive hits (average recall at 5% FDR) for CN-amplified genes after correction from 0.10 to 0.04 (*p* = 6.23 × 10^−29^) based on GISTIC [25] copy number scores from the CCLE, from 0.27 to 0.08 (*p* = 1.64 × 10^−8^) based on PicNic [26] copy number scores from the GDSC [13], and from 0.03 to 0.001 (*p* = 10^−4^) for non-expressed genes which are CN-amplified according to either GISTIC or PicNic scores. Additionally, true positive rates for known essential genes were slightly increased (average recall at 5% FDR) for *a priori* known essential genes from MSigDB [16] from 0.74 to 0.76 (*p* = 0.06), and significantly increased for essential genes from [15] from 0.59 to 0.63 (*p* = 8 × 10^−4^, **Supplementary Fig. S6**). The recall increment for known essential genes was greatest for lower quality CRISPR-KO data, suggesting that CRISPRcleanR contributes to a signal improvement in noisy or low quality data (**Supplementary Fig. S7**). Taken together, these results show that CRISPRcleanR is suitable for correcting bias in CRISPR-KO screening datasets generated with a variety of different sgRNA libraries.

### CRISPRcleanR preserves cell line essentiality profiles

We next determined whether the correction performed by CRISPRcleanR alters the overall essentiality profile of a given cell line. For Project Score data, we checked the position of sets of top-depleted sgRNAs from uncorrected logFCs along the profiles of corrected sgRNA logFCs by means of precision/recall analysis (**Fig. 4a, b**). We observed a median area under the precision/recall curve (AUPRC) of 0.92 (min = 0.81 for HCC-15, max = 0.96 for MDA-MB-436) for the top 50 depleted sgRNA, and a median AUPRC of 0.96 for the top 2,500 depleted sgRNA (min = 0.88 for HCC-15, max = 0.98 for MDA-MB-453). Considering that an experiment typically yields ∼6,000 sgRNAs called as significantly depleted with our library, this indicates that the CRISPRcleanR correction, while reducing false-positive rates, does not have an unwanted impact on the overall essentiality profile of a cell line.

**Fig. 4:**
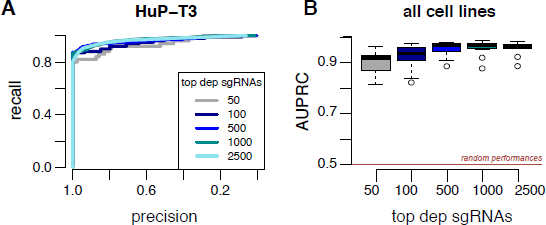
CRISPRcleanR retains overall essentiality profiles. (A) Example precision/recall curves in HuP-T3 cells for the indicated number of top depleted sgRNAs after CRISPRcleanR correction, classified based on their un-corrected sgRNAs logFC rank position. (B) Area under the precision/recall curves defined as for A for all cell lines. Box-plots show the median, inter-quartile ranges and 95% confidence intervals.

To further assess the impact of CRISPRcleanR on gene essentiality profiles, we compared all genes with a significant gain or loss-of-fitness effect before and after CRISPRcleanR correction as this is the key phenotype measured in CRISPR-KO screens (**Supplementary Fig. S8)**. For Project Score data, we found CRISPRcleanR impacted on the significant loss/gain-of-fitness effect for a median of 1.98% of all screened genes. This included a median of 24.69% genes significantly detected as exerting an effect on cellular fitness (gain- or loss-of-fitness) and a median of 17.02% of loss-of-fitness genes. The vast majority (88%) of these attenuated loss-of-fitness genes were composed of putatively false positive hits, involving genes which are not expressed (FPKM < 0.05), located in CN-amplified segments, prior known non-essential, or genes with a weak loss-of-fitness effect when compared to the whole set of genes called as loss-of-fitness in the uncorrected data (average logFC over the 4th quartile). For a very small number of genes (median 0.02% of genes, n = 28 unique genes total) the post-correction fitness effect was opposite to that observed prior to the correction. A very similar effect on significant genes following CRISPRcleanR correction was observed for the Project Achilles data (**Supplementary Fig. S8**). Thus, CRISPRcleanR preserves the overall essentiality profile present in a cell line and alters the significant fitness effects observed in the uncorrected data for only a minority of genes. Where correction occurs, the majority of instances involve likely putative false positive genes.

### CRISPRcleanR corrects sgRNA counts to enable mean-variance modeling

MAGeCK is a widely used computational tool to call gene depletion or enrichment in CRISPR-KO screens and is based on mean-variance modelling of median-ratio normalised sgRNA read-counts [22]. To make CRISPRcleanR compatible with mean-variance modeling approaches such as MAGeCK, we designed an inverse transformation to derive corrected sgRNA treatment counts from CRISPRcleanR corrected sgRNA logFC values. To benchmark our transformation, we compared results obtained from executing MAGeCK using normalised uncorrected and CRISPRcleanR corrected sgRNA counts by means of recall estimation when classifying predefined gene sets. The inverse transformation had an effect on both the mean and variance of the sgRNA counts, with the greatest impact on sgRNAs targeting genes in CN-amplified regions, whose value was consistently shifted toward the corresponding value in the plasmid/control condition (**Fig. 5a, b**).Furthermore, we observed a strong reduction in recall when classifying sgRNAs targeting genes in biased regions (PicNic scores ≥ 8 or GISTIC ≥ 2), when considering as positive predictions the sgRNAs called significantly depleted by MAGeCK. The median reduction was 75% for CN-amplified genes and 80% for CN-amplified non-expressed genes at a 10% FDR, and 72% and 100% reductions at a 5% FDR (**Fig. 5c** and **Supplementary Table S4**). In contrast, the effect on the recall of FE and non-essential genes was negligible (median = 2.9% reduction) (**Fig. 5c**). Thus, the reverse transformation post-correction enables the use of mean-variance modelling approaches such as MAGeCK for downstream calling of significant depletion or enrichment of genes.

**Fig. 5:**
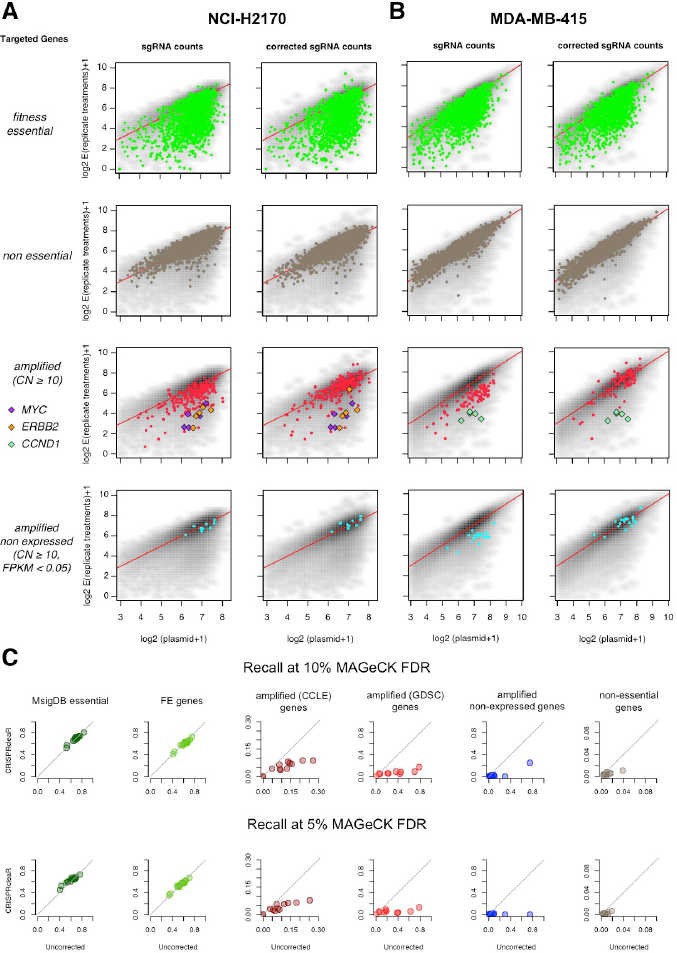
CRISPRcleanR corrected sgRNA counts and downstream analysis with MAGeCK. (A and B) Normalised counts of sgRNAs of the transfected libraries versus the control plasmid for FE and non-essential genes (first two rows of plots), CN amplified genes (third row) and CN non-expressed genes (fourth row), for two example cell lines before (first and third column) and after (second and fourth column) CRISPRcleanR correction. Essentialities for CN-amplified cancer driver genes such as *MYC*, *ERBB2* and *CCND1* are retained post correction. For the sake of readability only genes with at least 10 copies have been highlighted. (C) Comparison of recall using MAGeCK for sgRNAs targeting genes in six predefined gene sets when using as input CRISPRcleanR uncorrected and corrected sgRNAs counts.

### Robust detection of cancer dependencies following CRISPRcleanR

Since a major application of CRISPR-KO screens is the accurate identification of genes essential for cellular fitness in defined molecular settings, we investigated the ability of CRISPRcleanR to preserve the detection of expected cancer gene dependencies in individual cell lines. To perform a systematic analysis, we used CRISPRcleanR corrected sgRNA counts and a set of 64 cancer driver genes [27] which are modified by somatic mutation or CN amplification. We considered CN amplifications at the segment level (from [13]), thus including multiple genes in a segment.

Project Score cell lines included a total of 57 potential dependencies, involving a total of 29 cancer driver genes (9 mutated and 20 genes in amplified CN segments). Of these, we detected 21 dependencies prior to CRISPRcleanR correction (MAGeCK FDR < 10%), and 16 of them (76%) were preserved following CRISPRcleanR correction (**Supplementary Fig. S9 and Supplementary Table 5**). Examples included SW48 carrying the *EGFRg719s* mutation associated with depletion of *EGFR* targeting sgRNA, and MDA-MB-453 carrying the *PIK3CAh1047r* mutation associated with depletion of *PIK3CA* targeting sgRNA (**Fig. 6a**).

CRISPRcleanR preserved the ability to selectively detect cancer dependencies involving amplified cancer driver genes. For example, *MYC* is amplified in the cell line HT-29 and sgRNAs targeting *MYC,* as well as flanking genes, are reported as significantly depleted when using uncorrected logFCs (**Fig. 6b**). The logFC depletion is greater for *MYC* compared to other genes in this region. Following CRISPRcleanR correction, the sgRNAs targeting *MYC* remained significantly depleted, whereas those targeting the co-amplified flanking genes were no longer significant. A similar essentiality was selectively preserved post-CRISPRcleanR correction in an amplified region of chromosome 16 that contains *ERBB2* in the NCI-H2170 cell line (**Fig. 6c**). Two of the dependencies attenuated post correction involved co-amplification of two driver genes; *CDK12* co-amplified with *ERBB2* in NCI-H2170 and *CTTN* co-amplified with *CCND1* in MBA-MB-415 were no longer significant post correction. Similar results were found using the Project Achilles data with an overall retention rate of 80% (179 of 233) of dependencies post CRISPRcleanR correction (**Supplementary Fig S9 and Supplementary Table 5**). Of the attenuated dependencies, 41% (n = 44) involved genes co-amplified with another driver gene. In addition, we observed in both datasets a trend of increased significance (as measured by FDR) of detected dependencies post-correction. Overall, these results demonstrate that CRISPRcleanR allows for the accurate detection of cancer driver gene dependencies in CRISPR-KO datasets, including cancer genes residing within CN-amplified regions.

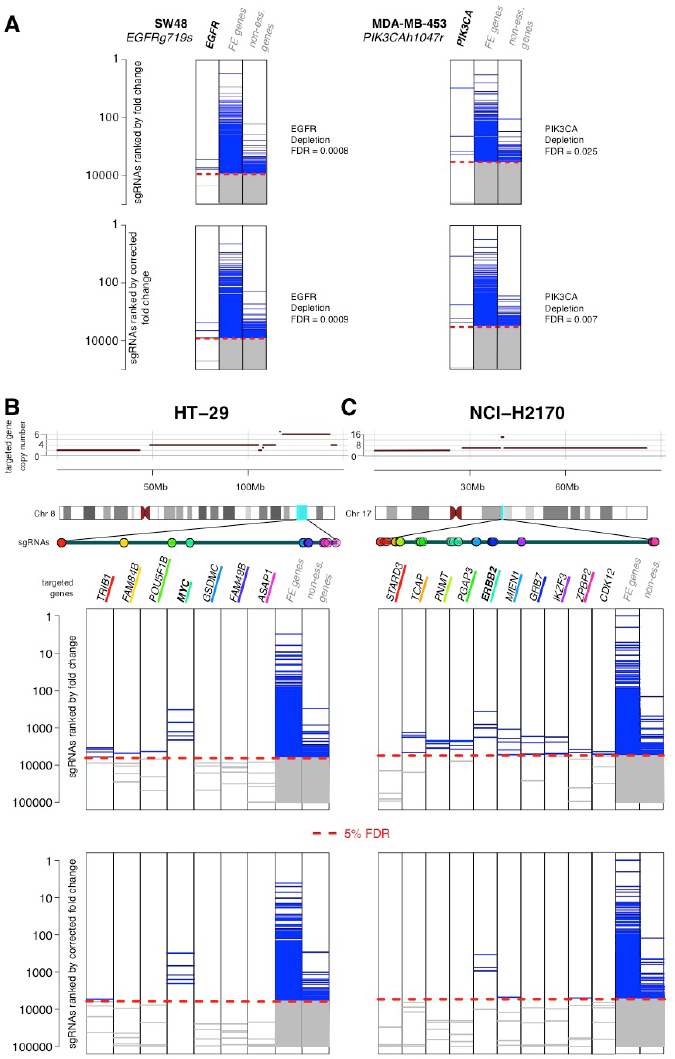

### Code availability and overview

CRISPRcleanR is implemented as an R [28] package and as an interactive Python package with full documentation, tutorials, built in datasets to reproduce the results in this manuscript, and is publically available (R package: https://github.com/francescojm/CRISPRcleanR and Python package: https://github.com/cancerit/pyCRISPRcleanR). The Python implementation is dockerized making it platform independent and usable in cloud environments (https://dockstore.org/containers/quay.io/wtsicgp/dockstore-pycrisprcleanr). CRISPRcleanR includes core functions for processing raw sgRNA count files for generating corrected sgRNA logFC values and corrected sgRNA counts for downstream analyses. CRISPRcleanR also includes functions to measure and visualise the extent and effect of the performed correction, the ability to detect CN-amplified non-expressed genes (which can be used as positive controls), and classification performances for *a priori* known sets of essential/non-essential genes pre- and post-correction.

## Discussion

In this study, we report CRISPRcleanR, a computational tool that detects genomic segments of gene-independent responses to CRISPR-KO in an unsupervised manner, and applies a segment-by-segment correction at the sgRNA-level for both fold-changes and read counts. The correction substantially reduces false-positive calls without altering true essentiality profiles and preserves known cancer gene dependencies within and outside of biased segments. CRISPRcleanR works on multiple genome-wide sgRNA libraries, and resulting corrected sgRNA logFC and read counts are compatible with downstream analyses performed by methods such as BAGEL or MAGeCK to statistically assess screen hits. CRISPRcleanR works efficiently irrespective of the sample size of the analysed dataset, even in single sample experiments.

Our motivation for developing CRISPRcleanR came from the observation that biases in gene essentialities observed in CRISPR-KO screens did not always show a linear correlation to their CN status, although biased segments are frequently associated with CN alteration. Additionally, in most of the cell lines analysed, variation in the mean logFCs of segments with the same CN were often greater than those between segments with different CN. Some cell lines showed greater bias in segments with lower CN. We even identified multiple instances of discontinuous bias on sgRNA logFCs within a particular CN segment, and biased responses within segments that are not CN-amplified. These observations argue for the development of methods such as CRISPRcleanR, which are independent of CN values for the analysis of CRISPR-KO screening data, and indicate that biased responses are not solely due to the amount of DNA damage and may also be caused by additional factors, such as local genomic structural variation (as recently reported in [29]).

CRISPRcleanR detects biased segments using sgRNA-level logFC in an unsupervised manner, eliminating the requirement for cell line CN information. This simplifies the analysis and is advantageous when reliable CN information is not available for a cell line; for example, when using a newly derived cancer cell model. In addition, cancer genomes are dynamic and continuously evolving, causing genetic variation between different clones of the same cell line. Genetic drift may occur during prolonged *in vitro* cell culture, due to different growth conditions (e.g. media composition), or in response to selective pressure (e.g drug treatment) and genetic manipulation (e.g. gene-editing). Thus, the genomic heterogeneity of cancer cells, even within clones of the same cell line, may confound CN-based correction methods when relying on pre-existing CN data, and negatively impact identification of gene essentialities. Furthermore, the performance of different copy number calling algorithms is variable and depends on the underlying genomic data available, and as a result this can be a further confounding factor when using CN-based correction methods. CRISPRcleanR overcomes these limitations by effectively correcting for biases in CRISPR-KO screens without requiring additional information about the cell models screened, and without making assumptions about the underlying cause of bias.

## Conclusion

CRISPRcleanR is a flexible tool implemented as R and Python packages to correct gene-independent bias found in whole-genome CRISPR-KO screens in an unsupervised manner at a single sample level. CRISPRcleanR facilitates the analysis of CRISPR-KO screens in cancer cells to identify essential genes.

## Methods

### Plasmids, cell lines and reagents

Cells were maintained in culture media as indicated in **Supplementary Table S1** in a 5% CO_2_ humidified incubator at 37 °C. The plasmids used in this study were from the mutagenesis toolkit described in [6] and are available through Addgene (Cas9 – 68343; CRISPR sgRNA library – 67989). Plasmids were packaged using the Virapower (Invitrogen) system as per manufacturer’s instructions.

### Genome-wide mutant library and screen

Cells were first transduced with lentivirus carrying Cas9 in T75 flasks at ∼80% confluence in the presence of polybrene (8 µg/ml). The following day, lentiviral containing medium was replaced with complete medium. Blasticidin selection was started on day 4 post transduction at a concentration determined from a titration in the parental cell line. Cas9 activity was assessed following selection using the Cas9 functional assay as described in [6] and a cut-off of 80% activity was applied (median = 89% activity across all cell lines). Cas9-expressing cells were maintained in blasticidin prior to transduction with the sgRNA library. Transduction with sgRNA library was carried out at ∼80% confluency with 3.3 × 10^7^ cells in T150 or T525 (triple layer) flasks, depending on cell size and surface area required, in technical triplicates. Cells were transduced with a predetermined viral amount that gives rise to ∼30% transduction, measured by BFP expression by cytometry, to ensure approximately 1 viral particle entering each cell based on a Poisson distribution model. Based on these initial cell numbers and transduction efficiency, the coverage of the sgRNA library (i.e. the number of cells containing each sgRNA) in each replicate was 100x. Puromycin selection commenced at day 4 to select for cells that had successful lentiviral integration. Actual library transduction efficiency and puromycin selection was analysed using flow cytometry before and after puromycin selection, respectively. A minimum number of 5.0 × 10^7^ cells were maintained at all times to ensure library representation was maintained. The cells were harvested 14 days post transduction and dry pellets were stored at −80 ºC.

Extraction of genomic DNA, PCR amplification of sgRNAs and Illumina sequencing of sgRNAs were carried out as described previously [3,6]. The number of reads for each sgRNA was determined using a script developed in-house.

### Data pre-processing and availability

sgRNA counts from both Project Score and Project Achilles (downloaded from: https://depmap.org/ceres/) were normalised assembling one batch per cell line, including the read counts from the matching library plasmid and all final read counts replicates, with a median-ratio method [30] to adjust for the effect of library sizes and read count distributions, after filtering out sgRNAs with less than 30 reads in the plasmid. Depletions/enrichments for individual sgRNAs were quantified as log_2_ ratio between post library-transfection read-counts and library plasmid read-counts. Finally, sgRNAs were averaged across replicates. This was performed executing the ccr.NormfoldChanges function of the CRISPRcleanR R package.

### Transcriptional and copy number data

Genome-wide substitute reads with fragments per kilobase of exon per million reads mapped (FPKM) for the 15 cell lines considered in this study were derived from the dataset described in [31]. Genome-wide gene level copy number data, derived from PicNic analysis of Affymetrix SNP6 segmentation data (EGAS00001000978) for the cell lines in the Genomics of Drug Sensitivity 1,000 (GDSC1000) cancer cell line panel [13], were downloaded from the GDSC data portal (dataset version: July 4th 2016), http://www.cancerRxgene.org. This dataset is also available at ftp://ftp.sanger.ac.uk/pub/project/cancerrxgene/releases/release-6.0/Gene_level_CN.xlsx. For each gene, the minimum copy number of any genomic segment containing coding sequence was considered. Additionally, gene level Gistic [25] scores obtained by processing Affymetrix SNP array data in the Cancer Cell Line Encyclopaedia [32] repository were downloaded from cBioPortal [33] (http://www.cbioportal.org/study?id=cellline_ccle_broad#summary).

### Analysis of gene-independent responses in cancer cell lines

For each cell line, segments of equal CN were identified by using CN data from the GDSC data portal [13,14] (as detailed below), and assigned a mean-logFC value by averaging across all of the sgRNAs targeting a segment. A CN *bias starting point* was computed for each cell line as the copy number value *n* > 2 such that statistically significant differences, as quantified by a Welch’s t-test, were observable between the mean-logFCs of segments of *n* CNs and those of segments of 2 CN. A CN *bias critical point* was computed for each cell line as follows. For each CN value *n* = 3, …, *m*-1 (with *m* = maximal segment CN value observed in the cell line under consideration), two univariate linear models were fitted, considering segment CN values as observations of the independent variable and the corresponding average segment mean-logFCs as those of the dependent one. The first model *P(n)* was fitted using CN values in {2, …, *n*} and corresponding average segment mean-logFCs, while the second one *L(n)* was fitted using CN values in {*n*+1, …, *m*} and corresponding average segment mean-logFCs. The *bias critical point* was then defined as the value *n* providing the large absolute difference between the slopes of the corresponding fitted models *P(n)* and *L(n)*.

### Calling significantly depleted sgRNAs and genes based on log fold-changes

All sgRNA were ranked by average logFCs derived from screening an individual cell line. This ranked list was used to classify sgRNAs targeting genes from two gold-standard reference sets of
FE and non-essential genes [15,21]: from now the *essential-sgRNAs* (*E*) and the non-essential-sgRNAs (*N*). For each rank position *k*, a set of predictions *P*(*k*) = {*s* ∊ *E* ∪ *N* : ϱ (*s*)≤ *k*}, with *ϱ*(*s*) indicating the rank position of *s*, was assembled and corresponding Precision (or Positive Predicted Value, *PPV*(*k*)) was computed as:

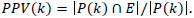

Subsequently the largest rank position *k*^*^ corresponding to a 0.95 Precision (equivalent to a False Discovery Rate (FDR) = 0.05) was determined as

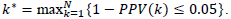

Finally, a 5% FDR logFCs threshold *F*^*^ was determined as the logFCs of the sgRNAs *s* such that *k(s)*=*k*^*^, and all the sgRNAs of the entire library with a logFC < *F*^*^ were considered significantly depleted at this FDR level.

To call depletion significance at a gene level, the same procedure was followed but averaging logFCs of sgRNAs targeting the same gene prior to the analysis, and considering ranks and positive/negative sets of genes instead of sgRNAs.

For the follow up analyses on the effect of correcting sgRNA treatment counts (computed as detailed below) we used the test function of the MAGeCK python package, indicating *none* as the value of the parameter specifying the normalisation method to use prior to the analysis, as a median-ratio normalisation was already applied to the analysed count files prior CRISPRcleanR correction.

### Receiver Operating Characteristic analyses

Across the different analyses, standard ROC indicators were computed considering as prediction sets significantly depleted sgRNAs (or genes) at a fixed level of 5% FDR (computed as detailed in the previous section or output by the MAGeCK tool), or genome-wide profiles of essentiality (as ranked lists of sgRNA logFCs, in some instances averaged on a per gene basis) to compute overall indicator curves, and using different positive/negative control sets (detailed below). To this aim, we made use of functions included in the pROC R package [34].

For the positive controls, sets of *a priori* essential genes were assembled by downloading relevant gene signatures from the MSigDB [16] (**Supplementary Table S6**). A list of ribosomal protein gene was derived from [17]. The consensual signatures resulting from this curation are available as individual data objects in the CRISPRcleanR R package.

### Segmentation analysis and logFC correction

Genome-wide essentiality profiles in the form of lists of sgRNAs logFC were sorted according to the genomic coordinates of the individual sgRNAs (library annotation and coordinates derived from [6]) using the function ccr.logFCs2chromPos of the CRISPRcleanR R package. Then, a circular binary segmentation algorithm [18,19] was applied using the ccr.GWclean function of the CRISPRcleanR R package, with a significance threshold to accept change-points *p* = 0.01, 10,000 permutations for *p*-value computation, a minimal number of 2 markers per region, and making use of the function segment from the DNAcopy R package [20] with other parameters set to default values.

Subsequently, sgRNA included in a segment had their logFCs mean-centered (across that segment) if collectively targeting at least *n* = 3 different genes, without pre-filtering any essential gene (differently from the sliding window approach used in [9]). This correction assumes that the true signal of loss/gain-of-fitness effect exerted by knocking-out a CN amplified gene sums up to a possible gene-independent impact on cellular fitness induced by targeting with CRISPR-Cas9 the chromosomal segment where that gene resides. By subtracting the logFCs mean to the sgRNA in the same detected biased segment, the gene-independent effect is flattened letting true fitness signals emerge. The possibility of using a median-based centering as a more robust alternative when the data is particularly noisy and/or many outliers are present (verifiable through a preliminary inspection of the uncorrected logFCs), for example due to dysfunctional or especially toxic sgRNAs, is also present in the implementation of CRISPRcleanR.

The minimal number *n* of targeted genes that a biased segment should contain in order to be corrected was adaptively determined by executing different trials of segments’ detection and correction varying *n* ∈ {2,3,5,10} and excluding/not-excluding from the analysis sets of *a priori* known essential genes assembled from MSigDB (as detailed in the previous section), collectively the *filter set*. Removing the filter set from a reference set of the FE genes yielded a *test set*. Areas under the recall curve (AURCs) were then computed evaluating the classification performances using as positive controls the test set, CN amplified genes, and CN amplified non-expressed genes (determined for each cell line) were then computed, across each trial using targeting sgRNAs’ logFCs before/after correction. For each of the positive control sets, reduction of recall (recall) were computed by comparing AURCs obtained before/after CRISPRcleanR correction.

Results showed that *n* = 3 provided the largest reduction of recall (**Supplementary Fig. S5**) of CN amplified and CN amplified non-expressed genes, and the lowest reduction of recall of the test set. Most importantly, this was observed invariantly with respect to removing/not-removing the filter set prior the analysis. As a conclusion, all the corrections presented in this manuscript were executed with this setting (*n* = 3 and without pre-filtering any gene). CRISPRcleanR package uses these settings by default, although offering to the user the possibility of changing them.

### Comparison of results across different libraries

Data from the mutagenesis of the HT-29 cell lines with the Brunello and Whitehead libraries were downloaded from the supplementary material of [24] and processed as described in the section *Data pre-processing and availability*. Correction outcomes were computed as detailed in *Receiver Operating Characteristic analyses*.

### Correction of sgRNA counts

We derived CRISPRcleanR corrected treatment count values for individual experiment technical replicates from the corresponding CRISPRcleanR corrected sgRNAs’ logFCs. To this aim, for each individual sgRNA, we first compute a CRISPRcleanR corrected treatment count averaged-across-replicate (first 7 formulas below), then we computed corrected treatment counts for individual replicates from this averaged value partitioning it across replicates proportionally to original (uncorrected) count values.

Formally, for each individual single guide RNA, a corrected treatment count *t_i_* was computed observing that:

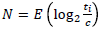

with *N* = CRISPRcleanR corrected logFCs for the sgRNA under consideration, *i* = 1, …, *n*, where *n* = number of treatment replicates, and *c* = counts of the sgRNA in the plasmid, and *E* indicates the mean function.

This implies

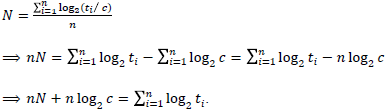

Assuming, for simplicity that all the *t*_*i*_ are the same (= *t*),

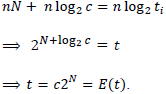

To derive the corrected counts for the individual replicates (which are obviously different from each other) from their mean, we keep constant the proportions seen in the uncorrected counts with respect to the sum of the counts across replicates:

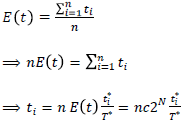

where 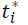 is the count of the sgRNA under consideration before correction in the *i*-th replicate and *T*^∗^ is their overall sum across replicates.

### CRISPRcleanR performances with respect to data quality

Cell lines from Project Achilles were grouped into 10 equidistant bins based on the quality of the corresponding profiles of gene essentiality, in increasing order. Data quality was quantified by the recall at 5% FDR for MSigDB [16] essential genes based on uncorrected fold-change rank positions. For each bin a variation of Recall pre/post-CRISPRcleanR correction was quantified, for 9 predefined gene sets, encompassing prior known essential/non-essential genes, copy number amplified genes and non expressed genes, as detailed in the previous sections.

### Evaluation of CRISPRcleanR correction on fitness gene calling

Gain/loss-of-fitness effect false discovery rate (FDR) scores were obtained by applying MAGeCK before/after CRISPRcleanR correction on the sgRNA counts from Project Score and Project Achilles. Percentages of attenuated fitness genes were computed as the ratio of genes with a significant gain/loss-of-fitness FDR (fitness genes), from the analysis of the uncorrected sgRNAs but not from the analysis of the corrected ones, with respect to the whole set of screened genes or the set of fitness genes detected in the uncorrected data, respectively. Percentages of distorted fitness genes were computed as the ratio of fitness genes detected in the uncorrected data which where still detected as fitness genes in the corrected data but with an opposite effect. Similar ratios were computed for attenuated/distorted loss-of-fitness and gain-of-fitness genes individually. The loss-of-fitness genes attenuated post-correction were further partitioned sequentially into the following disjoint sets across cell lines: non-expressed (with an FPKM < 0.05), copy number amplified (with a Gistic score > 1 or a PicNic copy number value > 2), prior-known non-essential (according to [15]), mild-phenotype (with a depletion logFC in the uncorrected data, averaged across targeting sgRNAs, falling over the 4th quartile of the logFCs of all the loss-of-fitness genes). Only cell lines with good quality data (recall for essential genes from [15] at 5% FDR > 0.5) and all data type (GISTIC and PicNic copy number, and basal expression FPKMs) available were included in this analysis.

### Retention of cancer driver gene dependencies following CRISPRcleanR correction

We performed a systematic unbiased case-by-case probing of putative oncogene addictions, by evaluating how corresponding dependencies are detected prior/post CRISPRcleanR correction, using data from Project Score and Project Achilles. From a list of 64 high confidence oncogenes [27], we considered those harbouring a cancer driver event (CDE), i.e. a cancer driver somatic mutation or a CN amplification as defined in [13], in at least one cell line of the two considered panels. For the Project Achilles, the analysis was restricted to 239 cell lines with genomic data available in [13]. The considered CN amplifications were at the chromosomal segment level and many of them included more than one oncogene. For each CDE observed in a given cell line, we then compared the loss/gain-of-fitness effect of the involved oncogene(s) observed prior/post-CRISPRcleanR in that cell line, quantified as MAGeCK FDRs. For Project Score, this resulted into 57 tested dependencies involving 29 CDEs (9 mutations and 20 CNAs encompassing multiple genes on the same segments). For Project Achilles, this resulted into 507 tested dependencies: 37 CDFEs (26 mutations and 11 CNAs encompassing multiple genes on the same segments).

## List of Abbreviations

AUPRC: Area Under the Precision/Recall Curve
AURC: Area Under the Overall Recall Curve
AUROC: Area Under the Receiver Operating Characteristic
CCLE: Cancer Cell Line Encyclopedia
CN: Copy Number
CRISPR-KO: CRISPR Knock-Out
DSB: Double Strand Breaks
FE: Fitness Essential
FPKM: Fragments Per Kilobase of Exon per Million reads Mapped
GDSC: Genomics of Drug Sensitivity in Cancer logFC Log Fold Change
logFC: Log Fold Change
MsigDB: Molecular Signature Database
ROC: Receiver Operating Characteristics
sgRNA: single guide RNA
TPR: True Positive Rate

## Declarations

### Ethics approval and consent to participate

Not applicable

### Consent for publication

Not applicable

### Availability of data and material

All data generated or analysed during this study are included in this published article and its supplementary information files. sgRNAs read count files analysed in this study are available for the HT-29, EPLC-272H, and A2058 cell lines as external data in the CRISPRcleanR R package, and they are used in the package vignette and documentation examples. sgRNA read counts for all cell lines used in this study have been deposited in BioStudies (https://www.ebi.ac.uk/biostudies/, Accession number S-BSST79).

### Author’s contributions

FI conceived the study, designed algorithms, processed data, performed validation tests, implemented and documented the R package, wrote and revised the manuscript; FMB contributed to the design of the study, performed screens and related experiments, curated data, contributed to documenting the R package, wrote and revised the manuscript; EG contributed ideas for the design of the study, curated data, revised the manuscript; SGB wrote python implementation of R package; EC pre-processed data from the Project Achilles; RS tested python implementation; CMB, RA, RP, PW, SH performed screens and related experiments; APB supervised the python implementation, ES contributed ideas for the design of the study, co-supervised it and revised the manuscript; JSR contributed ideas for the design of the study and supervised it, revised the manuscript; KY conceived the study and co-supervised it, designed the sgRNA library and the experimental setting of the screen, wrote and revised the manuscript; MJG conceived the study, designed the experimental setting of the screen, wrote and revised the manuscript, supervised the study.

## Acknowledgements

We thank David R. Wille, Vivek Iyer, Leopold Parts, Felicity Allen, Gabriele Picco, and Ultan McDermott for a number of insightful discussions. We thank Giuseppe Iorio for a number of acute and critical observations on the mathematics underlying this study.

## Additional Files

Additional File 1: Supplementary Fig.s S1 to S9 with legends and Legends of Supplementary Tables Additional File 2: Supplementary Tables S1 to S6

